# MLC Seq: *De novo* sequencing of full-length tRNA isoforms by mass ladder complementation

**DOI:** 10.1101/2021.05.22.445286

**Authors:** Xiaohong Yuan, Yue Su, Xudong Zhang, Spencer J. Turkel, Shundi Shi, Xuanting Wang, Eun-Jin Choi, Wenzhe Wu, Haichuan Liu, Rosa Viner, James J. Russo, Wenjia Li, Xiaoyong Bao, Qi Chen, Shenglong Zhang

**Affiliations:** Department of Biological and Chemical Sciences, New York Institute of Technology, New York, NY 10023, USA; Department of Computer Science, New York Institute of Technology, New York, NY 10023, USA; Division of Biomedical Sciences, School of Medicine, University of California, Riverside, Riverside, CA 92521, USA; Department of Chemical Engineering, Columbia University, New York, NY 10027, USA; Department of Pediatrics, University of Texas Medical Branch, Galveston, TX 77555, USA; Sealy Center for Molecular Medicine, University of Texas Medical Branch, Galveston, TX 77555, USA; the Institute of Translational Science, University of Texas Medical Branch, Galveston, TX 77555, USA; the Institute for Human Infections &Immunity, University of Texas Medical Branch, Galveston, TX 77555, USA; Thermo Fisher Scientific, 355 River Oaks Parkway, San Jose, CA 95134, USA

## Abstract

tRNAs can exist in distinct isoforms because of different chemical modifications, which confounds attempts to accurately sequence individual tRNA species using next generation sequencing approaches or to quantify different RNA modifications at specific sites on a tRNA strand. Herein, we develop a mass spectrometric (MS) ladder complementation sequencing (MLC-Seq), allowing for direct and simultaneous sequencing of full-length tRNA molecules, including those with low abundance. MLC-seq is achieved by improved instrumentation, and advanced algorithms that identify each tRNA species and related isoforms in an RNA mixture, and assemble full MS ladders from partial ladders with missing ladder components. Using MLC-Seq, we successfully obtained the sequence of tRNA-Phe from yeast and tRNA-Glu from mouse hepatocytes, and simultaneously revealed new tRNA isoforms derived from nucleotide modifications. Importantly, MLC-Seq pinpointed the location and stoichiometry changes of RNA modifications in tRNA-Glu upon the treatment of dealkylated enzyme AlkB, which confirmed its known enzymatic activity and suggested previously unidentified effects in RNA editing.

---

tRNAs have formidable complexities and are processed by multiple post-transcriptional regulatory mechanisms including base editing/modifications and addition of 3’ terminal bases^1,2^. Different tRNAs are responsible for transferring 20 amino acids and for decoding 61 different codons in mRNA^3^. tRNAs have isoacceptor and isodecoder forms, which may differ by as little as a single nucleotide or modification. Not every tRNA transcript copy is modified 100% of the time at each modification site. tRNA modifications are dynamic and subject to changes in cellular and external environments^4,5^. As such, tRNAs have many isoforms, and we do not even know how many tRNAs exist^6^. Currently there is no efficient way to resolve tRNA complexity^7^. Technically it has been difficult or even impossible to separate these isoforms for a specific tRNA species. It is also challenging for current next generation sequencing (NGS) methods to sequence them due to cDNA synthesis-blocking modifications. There are methods/enzymes used to remove these cDNA-blocking modifications^8^ that allow the tRNA’s primary sequences to be read, but at the expense of losing the modification information. Nanopore RNA-Seq has been used recently for sequencing of tRNAs^9^. However, it suffers from a high error rate and cannot reach single-nucleotide resolution for RNA modifications.

Mass spectrometry (MS) has been suggested as one of the most promising tools for studying RNA modifications in the field of epitranscriptomics^2,7,10^. However, MS alone has not been able to provide precise sequence and location information for these RNA modifications in complex biological samples. To obtain sequence information, many MS-based RNA mapping methods rely on other well-established complementary methods to supply such information, *e.g.,* NGS. With the prior sequence information of the sample determined, MS methods, especially tandem MS or MS/MS, have been used to confirm the specific sequences of interest and any related RNA modifications, based on mass shifts caused by nucleotide modifications (similar to peptide sequencing by MS^11^). However, compared to peptide fragmentation, fragmentation of RNA inside a mass spectrometer typically has more possible cleavage sites that generate a variety of potential fragment ions (a-w, b-x, c-y, d-z, and a-B), resulting in more complicated spectra and thus more difficulty in data interpretation (**Table 1**). Due to the spectral complexity, *de novo* sequencing of RNA by any MS/MS workflow could be challenging^12^. RNA mapping methods rely on matching MS/MS peaks of theoretical spectra predicted from a target sequence with the peaks of acquired spectra for analysis of nucleotide modifications. Significant recent developments of RNA mapping methods that provide better ion coverage by improved MS/MS fragmentation techniques, and increase data interpretation accuracy with better algorithms, have been reviewed^7,12,13,14^.

**Table. 1.**
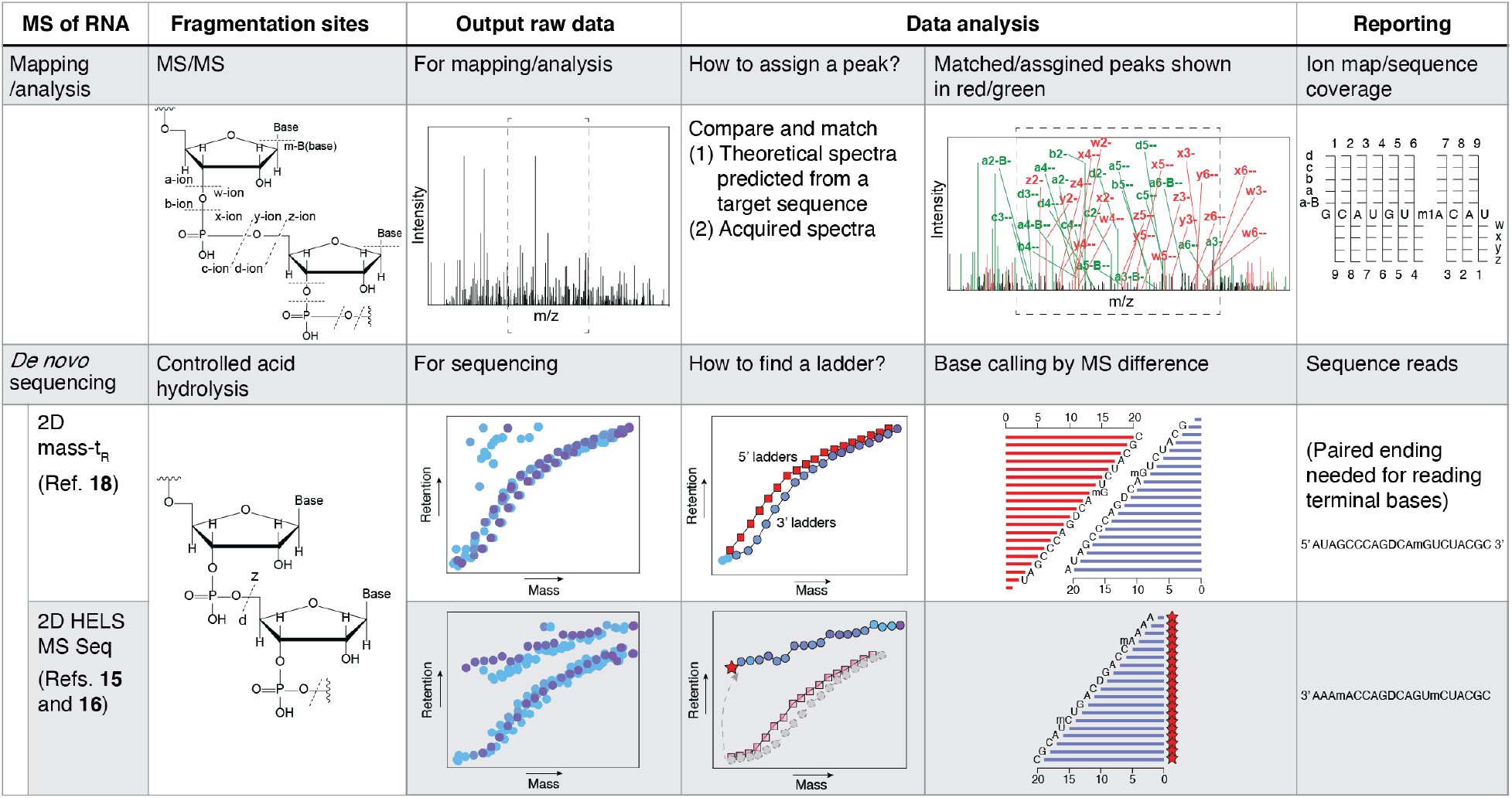
Comparison of previously reported MS-based RNA mapping/analysis and *de novo* sequencing. Due to its spectral complexity, *de novo* sequencing of RNA by any MS/MS workflow could be challenging. MS/MS-based RNA mapping methods must rely on an input sequence, *e.g.*, sequence information from other established sequencing methods, and matching of the theoretical spectra predicted from the targeted sequence with acquired spectra for sequence coverage analysis. However, MS-based *de novo* sequencing methods are able to report sequence reads without the need for any prior sequence knowledge because they rely on MS ladders generated by well-controlled acid hydrolysis, which form sigmoidal curves in 2D mass-retention time (t_R_) plots. 2D-Mass-t_R_ requires bi-directional paired-end reads for reading terminal bases (Ref. 18), while 2D-HELS MS Seq does not because the mass and t_R_ shift from a hydrophobic tag helps to identify/separate ladders (5’ and 3’ ladder) including terminal bases (Refs. 15 and 16).

Unlike MS-based RNA mapping methods, MS-based *de novo* sequencing methods rely on a complete set of MS ladders that are produced by exactly one random and unbiased cut on each RNA strand via controlled enzymatic or chemical degradation^7,10,15^. The ladder fragments carry nucleotide modifications from original parental RNAs, allowing identification, quantification and location of their nucleotide modifications^15,16^. However, each ladder must be perfect without any missing fragments in order to read all nucleotides in an RNA strand^15,16^. As such, they can provide *de novo* sequence information themselves, without the need for prior sequence information and thus are independent of other methods like NGS. However, it is difficult for all tRNA isoforms to meet this perfect mass ladder requirement, *e.g.*, due to variations in their abundances. We previously demonstrated the feasibility of using a MS laddering method for *de novo* sequencing of a mixture of 12 synthetic RNA strands with similar molar ratios, from which a perfect mass ladder was generated for each RNA strand without any missing ladder fragments^15^. This approach was also applied to sequencing of a yeast tRNA-Phe sample when coupled with a hydrophobic end-labeling strategy (HELS)^16^. However, only sequences of the major isoforms could be identified due to limited throughput from the sequencing algorithm which was tailored to report only the top-ranked sequences. Moreover, due to the short read length (<~35 nt per run), this method required T1 digestion to decrease the input tRNA length (~80 nt)^16^.

Here, we present a *de novo* mass ladder complementation sequencing method (MLC Seq) with significant enhancement in read length and throughput, making it possible to resolve tRNA isoform complexity (**Fig. 1**). MLC Seq allows direct and simultaneous sequencing of different tRNA isoforms from samples enriched for a specific tRNA, even those with low stoichiometry in a mixture (<1% relative abundance), in a single LC-MS run without perfect laddering. In reanalyzing a tRNA-Phe sample by MLC Seq, we discovered additional modifications and also the presence of tRNA-Tyr that was not identified previously. MLC Seq sequencing of tRNA-Glu obtained from mouse liver revealed new tRNA isoforms derived from nucleotide modifications. As proof of concept, MLC Seq then was used to track the location and stoichiometric changes of RNA modifications in the tRNA-Glu samples after enzymatic treatment, perhaps similar to what might occur during time-dependent enzymatic modification of *in vivo* samples. The ability to computationally separate MS ladder fragments and utilize imperfect MS ladders for sequencing is a paradigm shift for *de novo* MS sequencing of RNA, paving the way towards *de novo* MS sequencing of biological RNA at large scale. We expect this method will significantly expand the applications of MS-based sequencing to a broader variety of RNA samples, and provide a general sequencing tool for studying different RNA modifications in the field of epitranscriptomics.

**Fig. 1.**
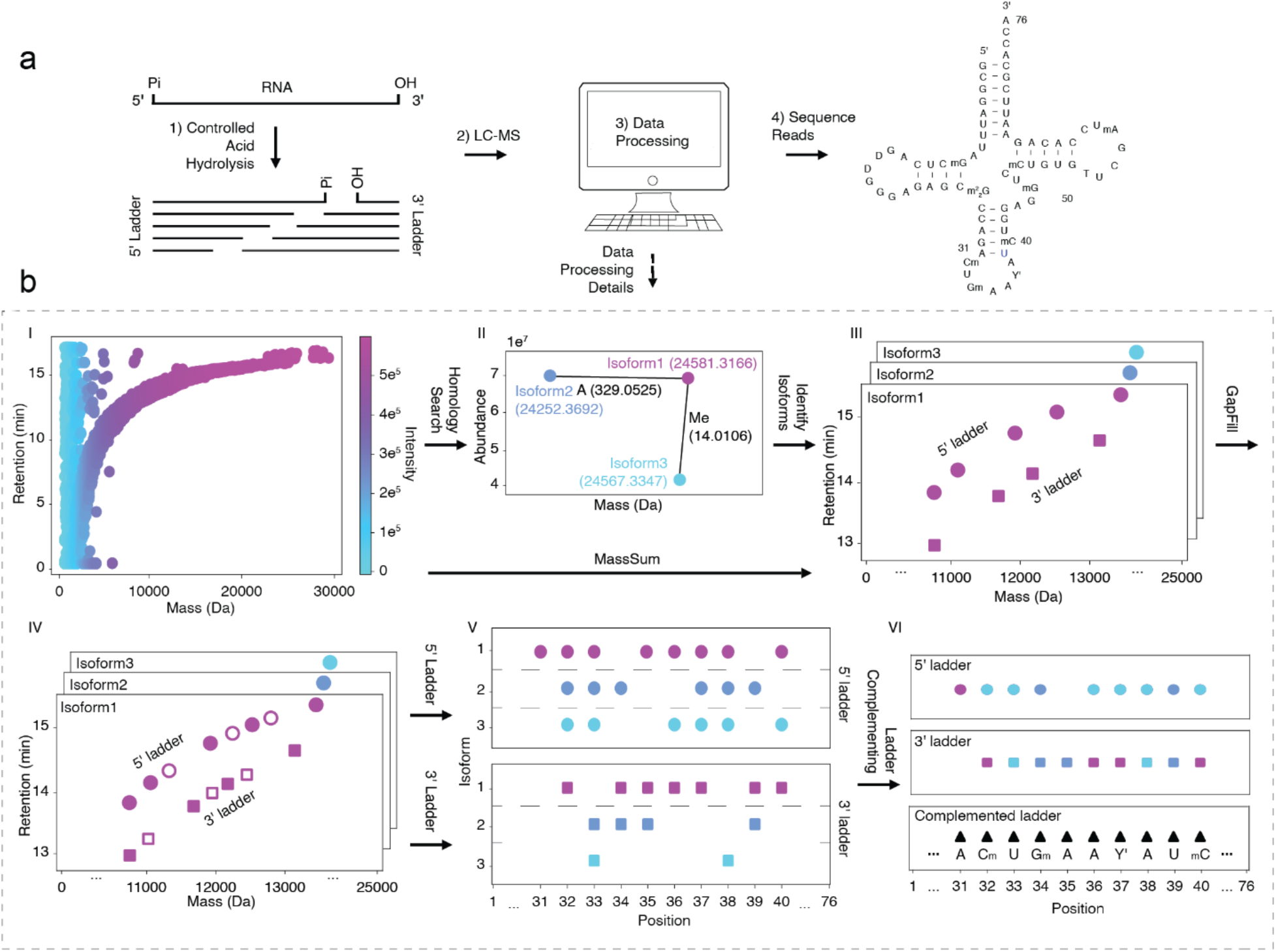
**a)** Workflow of *de novo* sequencing of tRNA isoform mixtures, including only four major steps: 1) acid hydrolysis of tRNA samples (single or mixed) under well-controlled conditions to generate ladder fragments, 2) LC-MS measurement of the resultant acid-degraded tRNA samples, containing tRNAs (intact or degraded) and all their acid-hydrolyzed fragments, and 3&4) data processing and generation of sequences consisting of both canonical and modified nucleotides. **b**) A complete set of step-wise computational tools, including algorithms for homology search, identifying acid-labile nucleotides, MassSum-based data separation, GapFill, ladder separation, ladder complementation, and sequence generation (see Methods).

## Results

### Direct sequencing of tRNAs together with modifications at single nucleotide resolution

We developed a MLC Seq method that can allow direct and *de novo* sequencing of full-length tRNAs without a cDNA intermediate (see workflow in **Fig. 1** and Methods). Using this method, the full-length 76 nt tRNA-Phe with all 11 modifications and full-length 75 nt tRNA-Glu with all 7 modifications as well as their isoforms were directly sequenced (**Fig. S1** and **Fig. S2**), without the T1 pre-fragmentation required by previous MS-based methods^16^. This MLC Seq method increased read length from ~35 nt to ~80 nt long per LC-MS run and sufficient throughput to identify and sequence each tRNA and related isoforms. This is achieved by 1) well-controlled acid hydrolysis to generate MS ladders; 2) use of the latest Orbitrap LC-MS (Thermo Fisher Scientific); 3) homology search of intact tRNAs to identify the related tRNA isoforms that share similar sequences but differ by as little as one nucleotide or modification (**Fig. 2a**), and to identify acid-labile nucleotides, *e.g.*, wybutosine (Y) at position 37 of the tRNA-Phe was converted to its depurinated ribose form (Y’) under acidic conditions (**Fig. 2b-c**); 4) development of strategies to identify and isolate MS ladders for each tRNA isoform in the mixture via computational tools of MassSum and GapFill (see Methods); and 5) MS ladder complementation combining imperfect ladders of different isoforms to form a complete MS ladder for sequencing. Since MS-based sequencing techniques rely on a unique mass value for identifying and locating each nucleotide, in the case where modifications have isomers with identical masses but different chemical structures such as pseudouridine (ψ) *versus* uridine (U) and methylation at different base positions, an extra step is required to differentiate these isomeric nucleotide modifications following our MS sequencing approach as described previously when sequencing of tRNA-Phe sample^16^. To differentiate different methylations in the tRNA-Glu, we included an additional step in the MLC Seq workflow by treating the tRNA-Glu sample with AlkB, which is a known enzyme that can demethylate tRNA-rich modifications such as 3-methylcytosine (m^3^C), 1-methyladenosine (m^1^A), and methylguanosine (m^1^G) (**Fig. S2**)^8^.

**Fig. 2.**
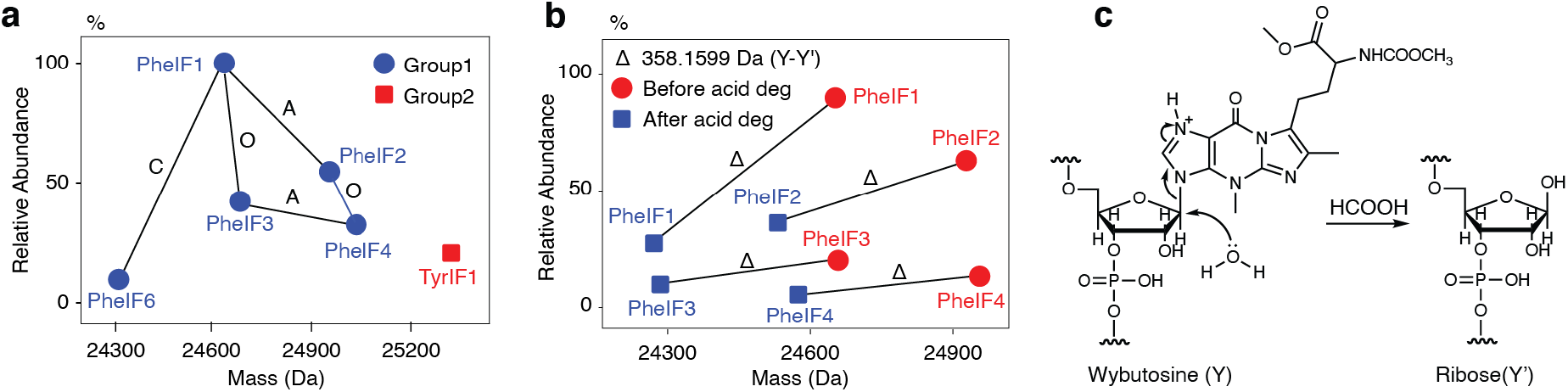
**a**) Homology search before acid degradation for identifying the related tRNA isoforms and cataloging them into different subgroups for MLC Seq. O is short for an oxygen, corresponding to a 15.9949 Da mass difference observed between different tRNA isoforms. **b**) Identification of each tRNA isoform containing acid-labile nucleotide modifications by comparing the mass changes of the intact tRNA before and after acid degradation. **c**) A mechanism illustrating a 358.1599 Da mass decrease due to the conversion of acid-labile wybutosine (Y) to its depurinated form (Y’) under acidic conditions.

### Identification and sequencing of all the isoforms

MLC Seq can begin to address the complexity of tRNA to reveal all the isoforms in a tRNA sample, especially for tRNA samples enriched for a specific tRNA species, but which may be a mixture of different isoforms caused by RNA editing, modifications and terminal truncations. This is achieved first by a homology search to identify each specific tRNA species and its related isoforms in a mixture at the intact, full-length tRNA level (**Fig. 2a**). For example, five tRNA-Phe isoforms were identified in the yeast tRNA-Phe sample (Sigma), and their stoichiometry was quantified (38.0%, 22.7%, 18.2%, 14.5%, and 6.5%, respectively) using their relative abundances together with their extracted ion currents (EIC)^15–17^. We randomly designated these isoforms as PheIF1, PheIF2, etc.; each has a distinct monoisotopic mass detected by MS (**Table S1**).

To allow MLC Seq method to sequence each related isoform in a mixture, we developed a new algorithm, designated MassSum, to identify and isolate ladder fragments related to each tRNA or isoform at the ladder fragment level. The MassSum strategy is based on the fact that the mass sum of any set of paired fragments (one in the 5’-ladder and the other in the 3’-ladder) generated during acid-mediated degradation of RNA by cleavage of one phosphodiester bond is constant (equivalent to the mass of each undegraded RNA plus the mass of a water molecule)^18^ (**Fig. 3a-b**). Since the mass sum is unique to each RNA sequence/strand, it can be used to computationally isolate MS data for all ladder fragments derived/degraded from the same tRNA isoform sequence in both the 5’- and 3’-ladders out of the complex MS data of sample mixtures containing multiple, distinct RNA strands (**Fig. 3c**).

**Fig.3.**
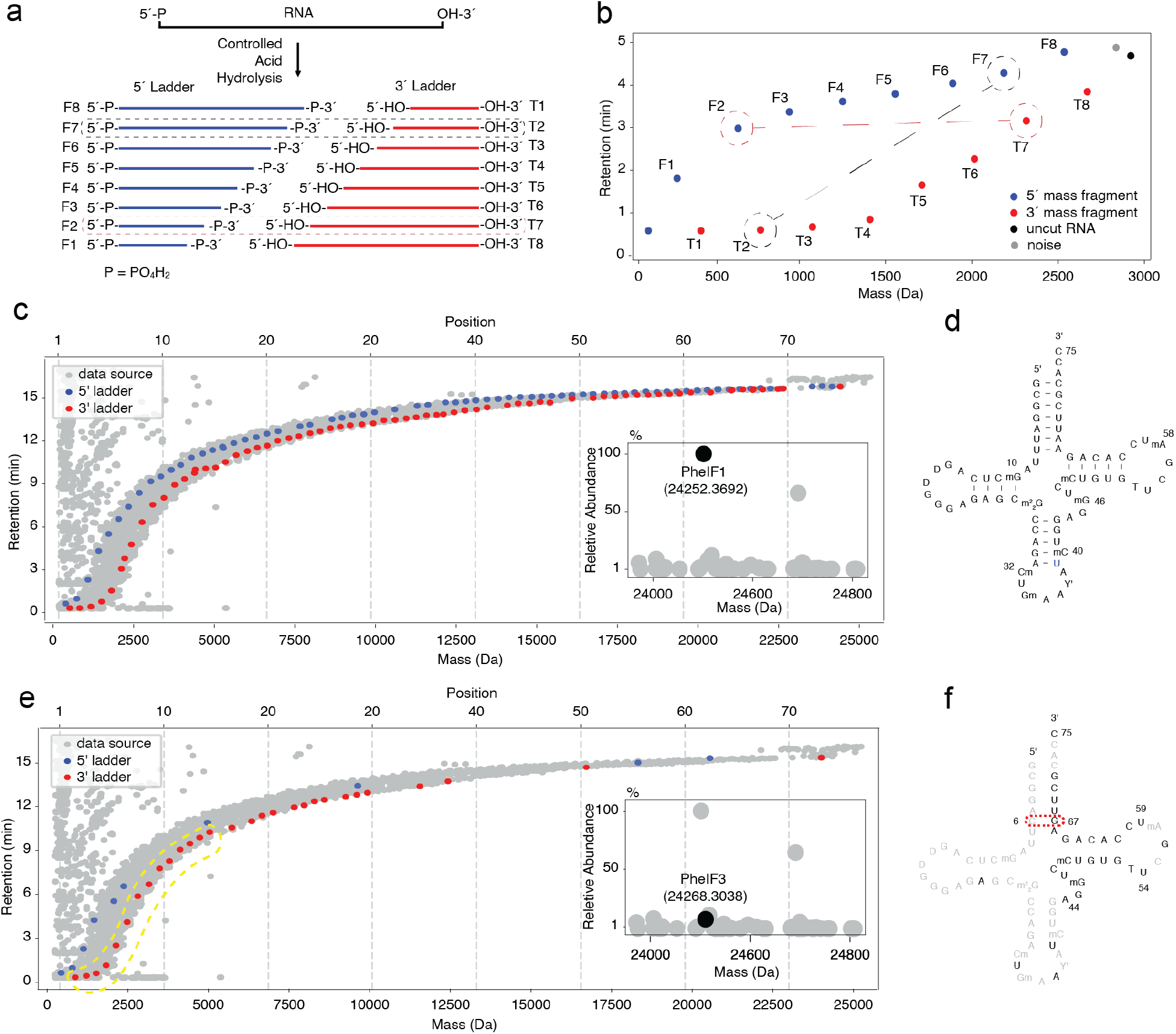
MassSum strategy and MassSum-based computational data separation. **a**) Isolated/mixed RNA starting material is partially digested in a manner that predominantly generates single-cut fragments. Taking a 9 nt RNA strand as an illustrative example, two ladder fragments are generated as a result of acid-mediated cleavage of the phosphodiester bond between the 1 st and 2nd nucleotide of the 9 nt RNA strand. One of them carries the original 5’-end of the RNA strand and has a newly-formed ribonucleotide 3’(2’)-monophosphate at its 3’-end (denoted F1). The other one carries the original 3’-end of the RNA strand and has a newly-formed hydroxyl at its 5’-end (denoted T8). **b**) The mass sum of any one-cut fragment pair, *e.g.,* mass sum of F2 and T7 or of F1 and T8, is constant and is equal to the mass of the 9 nt RNA plus the mass of a water molecule. Since the mass sum is unique to each RNA sequence/strand, it can be used to computationally separate all paired fragments of the RNA sequence/strand that were inextricable previously in the complex MS datasets. **c** and **e**) computational isolation of MS data for all ladder fragments derived/degraded from the same tRNA isoform sequence in both the 5’- and 3’-ladders out of the complex MS data of mixed samples with multiple distinct RNA strands. Separated data of 5’- and 3’-ladder fragments for 75 nt PheIF1 (major component of the sample mixture) (**c**) and and 76 nt PheIF3 (a tRNA-Phe isoform with C6 and G67) (~1% abundance compared to the PheIF1; minor component of the sample mixture) (**e**), respectively. **d, f**) *de novo* MS sequencing to generate the complete sequence of PheIF1 (**d**) and a partial sequence of PheIF3 (**f**), respectively.

With the MassSum-based data separation strategy, each tRNA isoform’s ladder fragments in the 5’- and 3’-ladders become identifiable *via* its unique individual intact mass, and each ladder fragment can be computationally separated out for MLC Seq sequencing by the MassSum algorithm. We identified and separated all ladder fragments related to five tRNA-Phe isoforms from MS data of the tRNA-Phe sample (**Fig. 3c** and **Fig. S1**). MassSum works even for the minor tRNA species in the complex RNA samples, such as PheIF3 which has only ~1% abundance compared to the PheIF1 (75 nt tRNA-Phe isoform with a monoisotopic mass of 24252.3692 Da after acid degradation) (**Fig. 3c-d**).

### Ladder complementation allowing sequencing of a broader range of tRNA samples

Traditionally, MS-based sequencing required a complete mass ladder with no missing ladder fragments in order to identify each nucleotide and its correct position in the sequence. However, very often a perfect ladder for a given tRNA isoform is not found after acid degradation and MS measurement, *e.g.,* due to its sample scarcity and/or the low stoichiometry of posttranscriptional modifications. Previously, such faulty ladders were considered dead ends in MS-based sequencing. Here with MLC Seq, we are able to fill in the imperfect ladder and thus resume the sequencing by combining the ladder fragments from other isoforms of the same tRNA group cataloged in the above-mentioned homology search.

Mass ladder complementation can be implemented within either the 5’- or 3’-ladder to contribute toward the completion of a perfect ladder with no missing ladder fragments. For example, when performing MLC Seq of the tRNA-Phe sample, the 5’-ladder fragment missing at position 3 of PheIF1 can be corrected site-specifically with the counterpart ladder fragment from PheIF2 (76 nt tRNA-Phe isoform with a monoisotopic mass of 24581.3166 after acid degradation) (**Fig. 4a**). Both isoforms have ladder fragments that can complement ladders for other tRNA-Phe isoforms. Even with the ladder complementation strategy, some ladder fragments in positions 1, 71 and 72 in the 5’-ladder are still missing (**Fig. 4a**). The missing ladder fragments at positions 32 and 34, however, are due to methylations on the 2’-hydroxyl group (Cm at position 32 and Gm at position 34) that block acid degradation^16,18^.

**Fig 4.**
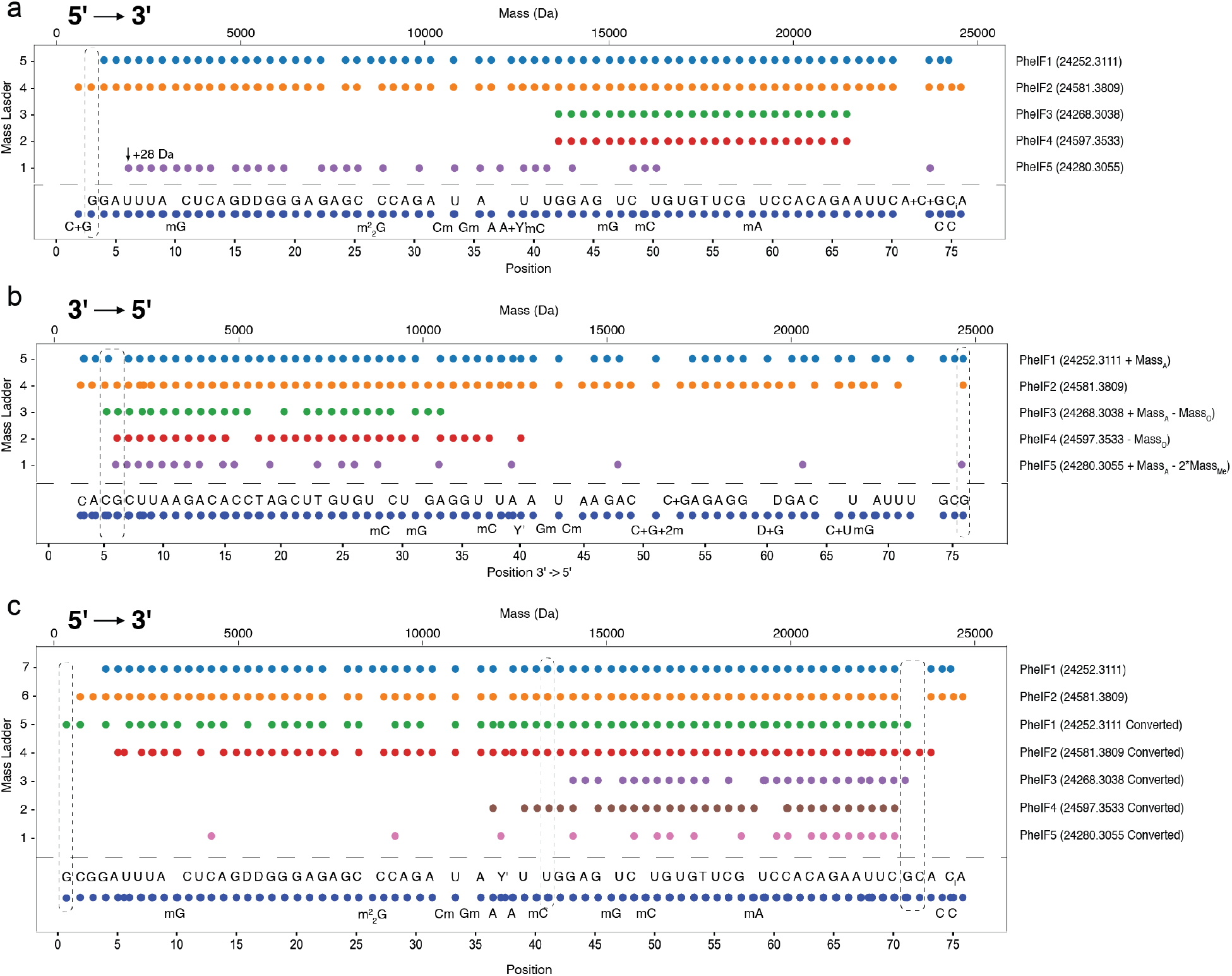
Perfection of a faulty mass ladder by complementing the missing ladder fragments from other isoforms identified through homology search. Ladder complementation can be implemented from different tRNA isoforms to perfect a 5’-ladder (**a**) and a 3’-ladder (**b**) of a specific tRNA-Phe species. **c**) All the 3’ ladder fragments can be converted to their corresponding 5’ ladder fragments for each tRNA isoform based on the MassSum principle. As such, each tRNA isoform could have two 5’-ladder fragments in each position on the 5’-ladder, one original 5’-ladder fragment and a second that was converted from its corresponding 3’ ladder fragment. These two 5’-ladder fragments can be used to re-affirm each other if both exist, or to complement each other if one is missing. This ladder complementation within the same tRNA isoform can be implemented together with cross-ladder complementation from different tRNA isoforms for sequencing of a specific tRNA. The missing ladder fragments at positions 32 and 34 are due to methylations on the 2’-hydroxyl group (Cm at position 32 and Gm at position 34) that block acid degradation. This would create a mass gap equivalent to 2 nucleotides, which was resolved previously by collision induced dissociation (CID) MS (Ref. 16).

To perfect the MS ladder for sequencing, the 3’-ladder can be used to correct the above missing fragments when sequencing the tRNA-Phe sample (**Fig. 4b**). Specifically, the 3’ ladder fragments at positions 76, 6 and 5 of PheIF2 can fill in the missing ladder fragments at positions 1, 71 and 72 in the 5’ ladder, respectively. After ladder complementation with the 3’-ladder, a perfect 5’-ladder having no missing ladder fragments can be formed for sequencing of all five tRNA-Phe isoforms (PheIF 1-5) in the tRNA-Phe sample (**Fig. 4c**).

### MLC Seq of different tRNA samples leading to new discoveries

Besides successful sequencing of three previously reported PheIFs in the Yeast tRNA-Phe sample (**Fig. 3** and **Fig. 4**)^16^, we also discovered other minor tRNA species that were not identified in this sample before, such as PheIF3 and tRNA-Tyr (both with <1% abundance compared to the most abundant PheIF1 in the sample). For example, two short sequences, T56-U62 and C66-A75, can be read and can be used to blast search the entire tRNA-Tyr sequence, *e.g.*, from reported tRNA sequences in literature/databases, which can be used, in turn, to find more ladder fragments for subsequent verification and modification analysis (**Fig. S3**).

To test our method’s applicability, we also sequenced a mouse liver tRNA-Glu sample that was purified by an affinity pulldown assay combined with gel recovery (**Fig. 5a**). Despite efforts attempting to obtain highly purified tRNA-Glu, we still found that multiple isoforms exist as a result of partial nucleotide modifications of the tRNA-Glu (**Fig. 5b**). The two most abundant tRNA-Glu isoforms (GluIF1 and GluIF2) have a 14.0081 Da mass difference, corresponding to a methyl group. The fact that these two isoforms with a 14.0081 Da mass difference (or within 10 PPM^19^) co-existed in the sample suggested that one of the nucleotides of the tRNA isoform was partially modified. With MLC Seq, we were able to sequence both tRNA-Glu isoforms (**Fig. 5c-e**). To our surprise, despite sharing the same primary sequence with tRNA-Glu, the tRNA-Glu has a very different nucleotide modification profile than what was reported before^20^. Specifically, we discovered a partially modified methylated A (which was predicted as m^1^A because it was demethylated 100% to A by AlkB, see **Fig. 5c-d**) at position 57 of the isoform (68% m^1^A vs. 32% A) and a partially methylated G (which was predicted as m^1^G because it was demethylated 88% to G by AlkB, see **Fig. 5c-d**) at position 10 of the isoform (48% m^2^G vs. 52% G). Also, instead of pseudo-U, we observed a dihydro-U (D) at position 20 in the isoform (**Fig. 5d-e**).

**Fig. 5.**
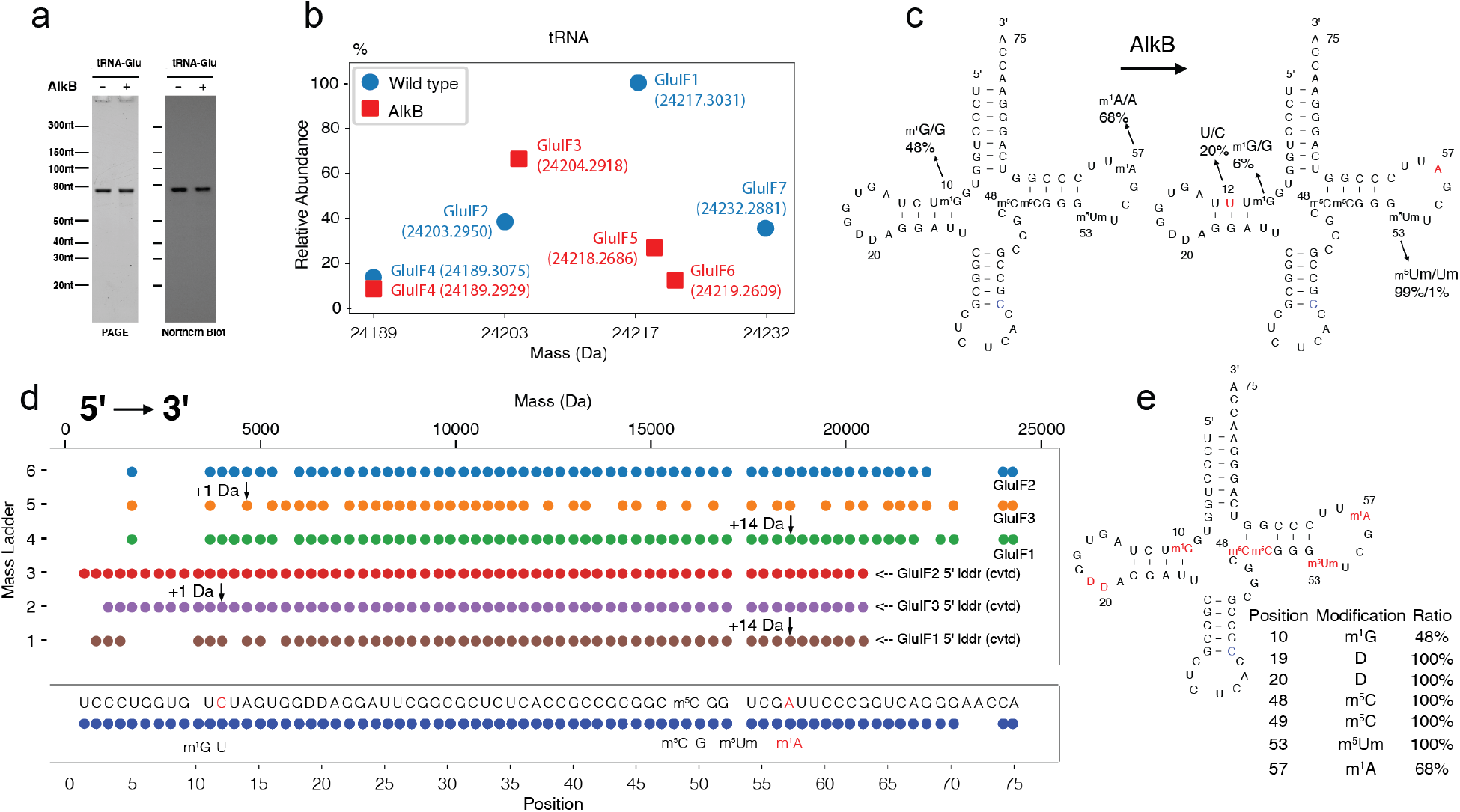
MCL Seq of the wild-type tRNA-Glu samples from a mouse liver and monitoring the location and stoichiometry change of nucleotides in the tRNA after AlkB-treatment experiments. **a**) Northern blot data on tRNA-Glu. PAGE gel image (left) and Northern blot (right) confirmation of enriched tRNA-Glu from mouse liver. **b**) Monoisotopic mass differences between wild-type and AlkB-treated intact tRNA-Glu isoforms. The mass difference, for instance, between the most abundant wild-type tRNA-Glu isoform GluIF1 and the most abundant AlkB-treated tRNA-Glu isoform GluIF2, is 13.011Da. Also, the ratio of methylated G (likely to be m^1^G) at position 10 was reduced from 48% to 6% after AlkB treatment. **c**) The 13.011 Da mass decrease caused by AlkB treatment can be tracked down to a demethylation of m^1^A at position 57 (yield: 100%) and a deamination of C to U at position 12 (20%). The 28.024 Da mass decrease from GluIF7 to GluIF3 is likely due to losing 2 methyl groups, one demethylation of m^1^A at position 57 (yield:100%) and m^1^G at position 10 (yield: 88%). About 1% of m^5^Um in position 53 was demethylated to Um after AlkB treatment (See Table S2). **d**) Mass ladder complementation during MCL Seq. The missing ladder fragment at position 53 is due to a methylation on the 2’-hydroxyl group of m^5^Um that blocks acid degradation. **e**) MLC Seq reveals different isoforms, base modifications and base editing, as well as their stoichiometries, in the tRNA-Glu sample from a mouse liver. The assignment of m^1^G at position 10, m^1^A at position 57, and m^5^C at positions 48 and 49 over other methylated isoforms was made largely by reported AlkB activities in reference to known nucleotide modifications reported in tRNA-Glu.

### Monitoring dynamic changes of nucleotides/modifications in tRNA

Due to their wealth of modifications, it is difficult to sequence tRNAs, and even more challenging to track their dynamic changes. To test our method’s capacity to determine dynamic nucleotide modifications changes, we treated the tRNA-Glu mouse hepatocyte sample with AlkB, which was also used to differentiate isomeric methylations in the tRNA-Glu. After AlkB treatment of the tRNA-Glu sample, GluIF1 disappeared, and another isoform GluIF3 with a 13.011 Da less mass became the most abundant (**Fig. 5b**). The MLC Seq results indicated that the 13.011 Da mass decrease was caused by a demethylation of m^1^A at position 57 (100% yield) and a C to U deamination at position 12 (20%) (**Fig. 5c-d**). GluIP3 may also be generated by removal of two methyl groups from GluIF7, one demethylation of m^1^A at position 57 (yield:100%) and m^1^G at position 10 (yield: 88%). In fact, we found that the ratio of m^1^G at position 10 was reduced from 48% to 6% after AlkB treatment, showing that the m^1^G is sensitive to AlkB, which was also confirmed by a dramatic intensity decrease of m^1^G-containing MS ladder fragments beginning at this position (**Table S2**). These results suggest that the nucleotide modification at position 10 might be different than the m^2^G reported before^20^ and might be a m^1^G instead, which can be partially demethylated by AlkB. To our surprise, we found that about 1% of 5,2’-O-dimethyluridine (m^5^Um) in position 53 was demethylated to 2’-O-dimethyluridine (Um) after AlkB treatment (See **Table S2**). The minor tRNA-Glu isoform GluIF7 also produced GluIF5 after demethylation of m^1^A at position 57, resulting in a 14.0195 Da mass decrease.

## Discussion

MLC Seq is a significant methodological advance because it results in direct and *de novo* sequencing of full-length tRNA without cDNA conversion, and simultaneously reads out all the nucleotide modifications at single-nucleotide resolution. Algorithms in MLC Seq can incorporate partial ladders with missing fragments, identify each RNA in a mixture, and computationally separate all MS ladder fragments for sequencing of each RNA isoform, including even tRNA isoforms with low stoichiometry, from the RNA mixture. Of practical significance is its ability to resolve RNA sample complexity, such as different tRNA isoforms, and to use LC-MS to sequence RNAs with even faulty or incomplete mass ladders. This ability greatly expands the method’s applicability, allowing sequencing of a broader range of RNA samples that cannot otherwise generate perfect MS ladders. These problematic samples include those with tRNA in low abundance and/or with rare modifications. When MLC Seq *de novo* sequencing was applied to a previously reported tRNA-Phe sample, we discovered additional tRNA-Phe isoforms, *e.g.,* A to G transition at position 67 of PheIF1 and PheIF2, and also the presence of tRNA-Tyr in low stoichiometry, that were not identified previously, thereby confirming the ability of the method to uncover greater complexities of tRNA than previous methods.

The ability of MLC Seq to track the complexity of tRNA has significance for biological studies. For example, we discovered new tRNA isoforms derived from nucleotide modifications such as m^1^A and D, which calls for further investigation. Because RNA modifications are dynamic and subject to change, of particular importance for biological studies is the ability to track changes of nucleotides in tRNA as a result of cellular activities. We used MLC Seq to analyze wild-type and AlkB-treated tRNA-Glu samples obtained from mouse liver. MLC Seq was used to pinpoint the locations and stoichiometric changes of tRNA modifications, *e.g.*, due to demethylation and deamination, before and after treatment with the dealkylating enzyme AlkB. The sequence information is the first detailed description of the tRNA-Glu, which includes two back-to-back dihydro-Us (D) at positions 19 and 20, and likely a partially modified m^1^A at position 57 and a partially modified m^1^G at position 10. In addition, the m^5^Um in position 53 of tRNA-Glu was found to be sensitive to AlkB treatment. It is expected that the new information will provide further insight in understanding the biological functions, *e.g.*, stability of the parental tRNAs and the resulting cleavage of tsRNAs, which has implications in neurodevelopmental disorders, metabolic disorders, and infectious diseases^21–28^.

Many efforts have been made to improve MS/MS or MS^n^, *e.g.*, for analysis of small metabolites and peptides/proteins^11^. If similar improvements could be made for monoisotopic mass measurements of RNA, including better instrumentation and data processing software, we can be very optimistic about the future of MS-based sequencing of RNA. The full potential of the method’s sequencing read length and throughput remains to be explored, and it seems instrument-dependent, *i.e.,* mass spectrometers with higher resolving powers and better sensitivity may lead to increased read length and throughput, and lower sample requirements. With more advanced LC-MS instruments, we expect the read length can be increased to much longer than ~80 nt per run, allowing direct sequencing of RNAs longer than tRNAs such as rRNA and viral RNA together with their nucleotide modifications. This paves the way for large scale *de novo* MS sequencing of complex biological samples, especially with state-of-the-art LC-MS instruments and their built-in capacity for automation. Thus, this MLC Seq method will provide a general sequencing tool for studying RNA modifications, which is urgently needed more than ever, considering for example that >40 unidentified nucleotide modifications have been reported in SARS-CoV-2 RNA^29^, though their identities and functions remain unknown. Such a method will be instructive for studying SARS-CoV-2 RNA and other RNAs and to unravel epitranscriptomic roles in COVID-19 and other diseases.

## Methods

### Workflow of *de novo* sequencing of tRNA isoform mixtures

The workflow includes four major steps (**Fig. 1**): 1) acid hydrolysis of tRNA samples (single or mixed RNAs) with well-controlled conditions to general 3’ and 5’ ladder fragments^1–4^, 2) LC-MS measurement of the resultant acid-degraded tRNA samples, containing tRNAs (intact or degraded) and all their acid-hydrolyzed fragments, 3&4) data processing, including identification and separation of 3’ and/or 5’ MS ladder fragments thereby generating the sequence of one or more RNA molecules and detecting the presence, identity, location, and quantity of any RNA nucleotide modifications.

### Acid hydrolysis degradation of tRNA

Formic acid was applied to degrade tRNA samples, including tRNA-Phe sample (Sigma) and cellular tRNA-Glu samples (see Sections of tRNA-Glu sample preparation), for producing mass ladders, according to reported experimental protocols^1–4^. In brief, we divided each RNA sample solution into three equal aliquots for formic acid degradation using 50% (v/v) formic acid at 40 °C, with one reaction running for 2 min, one for 5 min and one for 15 min. The reaction mixture was immediately frozen on dry ice followed by lyophilization to dryness, which was typically completed within 30 min. The dried samples were combined and suspended in 20 μL nuclease-free, deionized water for LC-MS measurement.

### Liquid chromatography-mass spectrometry (LC-MS) analysis

The acid-hydrolyzed tRNA samples were separated and analyzed on an Orbitrap Exploris 240 mass spectrometer coupled to a reversed-phase ion-pair liquid chromatography (ThermoFisher Scientific, USA) using 200mM HFIP and 10mM DIPEA as eluent A, and methanol, 7.5 mM HFIP, and 3.75mM DIPEA as eluent B. A gradient of 2% to 38% B over 15 minutes was used to elute RNA samples across a 2.1 x 50 mm DNAPac reversed-phase column. The flow rate was 0.4 mL/min, and all separations were performed with the column temperature maintained at 40 °C. Injection volumes were 5-25 μL, and sample amounts were 20-200 pmol of tRNA. tRNAs were analyzed in a negative ion full MS mode from 410 m/z to 3200 m/z with a scan rate of 2 spectra/s at 120k resolution. The sample data was processed using the Thermo BioPharma Finder 4.0 (ThermoFisher Scientific, USA), and a workflow of compound detection with deconvolution algorithm was used to extract relevant spectral and chromatographic information from the LC-MS experiments as described previously^1–4^.

### Homology search

Once LC-MS data is output into a 2D mass-retention time (t_R_) plot, we start a homology search of intact tRNAs in the mass range of >~24k Dalton (or ~75 nt; on average ~318 Dalton/nt) using an in-house developed algorithm (See **Fig. 2a**) to first identify related tRNA isoforms that may share the same ancestral precursor tRNA, but are different, *e.g.,* in posttranscriptional profiles of nucleotide modifications, editing, and truncations. Mass differences between two intact tRNA isoforms are calculated and matched to the known mass of each nucleotide or nucleotide modification in the database^4^. For example, known mass difference between these intact tRNAs such as 14.0157 Da and 329.0525 Da (with PPM difference <10 ppm)^5^ can be assigned to a methylation (Me/-CH2-) event and an additional A nucleotide, respectively. Therefore, these intact tRNAs are assigned to the same tRNA group and considered as homologous isoforms of a specific tRNA whose sequences are decoded together.

### Identification of acid-labile nucleotides

Acid-labile nucleotides are identified using another computational tool implemented in Python. The tool analyzes the connections between the compounds before acid degradation and the ones after acid degradation. For each such compound pair, if the monoisotopic mass difference can match a mass difference calculated from the possible structural change to a specific nucleotide modification during acid hydrolysis, or match the mass difference sum of a subset of different acid-labile nucleotide modifications, the compound pair would be selected and further considered as potentially containing acid-labile nucleotide modifications. In general, if the intact mass of an RNA species did not change after acid degradation, this intact mass will be used for MassSum data separation. If the intact mass did change after acid degradation, an acid-labile nucleotide may be identified by matching the observed mass difference with the theoretical mass difference caused by acid-mediated structural changes of the nucleotide (See **Fig. 2b-c**).

### MassSum data separation

MassSum is an algorithm developed based upon the acid degradation principle presented in **Fig.3**. Taking advantage of the fact that each fragmented pair from two ladder groups (5’ and 3’ ladders) sums to a constant mass value that is unique to each specific tRNA isoform/species, the algorithm can isolate ladder compounds corresponding to a specific tRNA isoform. MassSum simplifies the dataset by grouping MS ladder components into subsets for each tRNA isoform/species based on its unique intact mass. Since the well-controlled acid degradation reaction cleaves RNA oligonucleotides at one specific site of the phosphodiester bond, on average, one cut per RNA^4^, the masses of two RNA fragments (Mass_3’ portion_ and Mass_5’ portion_) from the same strand add up to a constant value (Mass _sum_).

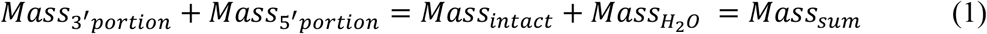

Taking advantage of this unique mass sum between the paired 3’ and 5’-ladder fragments (*Equation 1*), the algorithm chooses two random compounds from the acid-degraded LC-MS dataset and adds their mass values together, one pair at a time. If the sum of the two selected compounds is equal to a specific Mass_sum_, these two compounds will be selected into the pool for each RNA accordingly. The process repeats until all compound pairs have been inspected. In the end, MassSum will cluster the dataset into different groups; each group is a subset that contains 3’ and 5’-ladders of one RNA sequence.

### GapFill

GapFill is another algorithm developed as a complement to MassSum. Since MassSum handles compounds in pairs, in the case that one ladder fragment is missing, *e.g.,* in the 5’-ladder, the corresponding single-cut ladder fragment, even if it exists in the 3’-ladder, will not get separated/called out by the MassSum algorithm. In order to pull out all the ladder fragments from the complex MS data, a GapFill algorithm was designed to rescue any ladder fragments missed by MassSum separation. We identify a gap that has ladder fragments missing between compounds, *e.g.*, Mass_5’i_ and Mass_5’j_ in the 5’-ladder. Among the gaps there exist many compounds in the degraded LC-MS dataset, but none were selected out after MassSum algorithm during data separation. GapFill iterates over each potential compound in the gap in the original LC-MS dataset, examines the mass differences of this compound and the two ending compounds with Mass_5’i_ and Mass_5’j_. If the mass difference is equal to the sum of one or more nucleotides or modifications in the RNA modification database^4^, we define it as a connection. If the compound in the gap has connections with both ending compound Mass_5’i_ and Mass_5j_, this compound would be selected into a candidate pool for the later sequencing process. After iteration, GapFill calculates connections of the compounds in the candidate pool and assigns weights to them based on the frequency of each connection. The compounds that contain the highest weights would be the ones chosen to fill in the gap.

### 5’- and 3’-Ladder separation

The t_R_ differences can be used to further computationally separate these two ladders, breaking two adjacent sigmoidal curves into two isolated curves, one for the 3’- and the other for the 5’-ladder (**Fig. 1b**). Due to a large amount of RNA/fragment compounds, the dividing line between two subsets of 5’- and 3’-ladder fragments is not visually decisive in the 2D plot. Thus, we developed a computational tool to separate the 5’ and 3’ fragments. We roughly divide all the compounds in each LC-MS data pool into two subgroup areas by circling compounds in the top collective curve of the 2D mass-t_R_ plot and marking the compounds as 5’-ladder fragment compounds, while the compounds in the bottom curve are marked as 3’-ladder fragment compounds. The purpose of selecting the top area is to include as many 5’ fragment compounds as possible, in the meanshile few 3’ fragments as well. Similarly, the purpose of selecting the bottom area is to include as many 3’ fragment compounds as possible, with few 5’ fragments. Overlap between two selected ladder subgroups is inevitable, due to limited t_R_ differences between these two subgroups. The aim of the manual selection step is not to separate the 5’ and 3’ fragments with high precision, but to serve as two input ladder fragments for an algorithm to output 5’ and 3’ ladder fragments separately for each tRNA isoform/species.

### Ladder complementation and generation of RNA sequences

After MassSum and GapFill, each tRNA isoform has its own set of separate 5’-and 3’-ladders (not combined). Each ladder (5’- or 3’-) consists of a ladder sequence, and we can determine if these ladders are perfect without missing any ladder fragment so as to read the first to the last nucleotide in the RNA sequence. If not, we can complement ladders from other related isoforms in order to get a more complete ladder for sequencing. A computational tool was designed to align these ladders based on the position from the 5’→3’ direction. For example, we layout each tRNA-Phe isoform’s ladder, *e.g.*, 5’-ladder in **Fig. 4a**, on top to each other vertically; horizontally we align the 5’-ladder of each isoform according to the position of each corresponding ladder fragment, ranging from position 1 to 76 nt for tRNA-Phe (~318 Dalton/nt). Ladder complementation can be performed separately on 5’ or 3’ ladders, resulting in one final 5’ ladder or one final 3’ ladder separately. Additionally, all the 3’ ladder fragments can be converted to their corresponding 5’ladder fragments for each tRNA isoform based on the MassSum principle. As such, each tRNA isoform could have two 5’-ladder fragments in each position on the 5’-ladder, one original 5’-ladder fragment, and a second that was converted from its corresponding 3’ ladder fragment, for reaffirmation and/or complementation.

### Validation and confirmation of RNA sequence reads using anchor-based sequencing algorithm

To validate and confirm the RNA sequence reads, we use an anchor-based sequencing algorithm^1^ to read out the RNA sequence directly from perfected mass ladders that are obtained from the previous step. There are three main steps in the Anchor-based Sequencing Algorithm: (1)

Anchor-based base calling, which detects and outputs all the canonical and modified nucleotides starting from the anchor node; (2) Depth-First Search (DFS)-based draft sequence reads generation, which connects the adjacent canonical and modified nucleotides together and outputs them as draft sequence reads; and (3) final sequence identification based on the global hierarchical ranking strategy (GHRS), in which the draft sequence reads will be ranked according to a set of ordered criteria, such as the number of canonical and modified nucleotides (a.k.a, read length), average abundances/intensities, and average PPM.

### Mouse liver tRNA-Glu pulldown

The tRNA-Glu was purified by affinity pulldown assay combined with gel recovery, with modified protocols from the previous report^6^. The total RNA of mouse liver was harvest by TRIzol^TM^ reagent (Invitrogen^TM^ 15596026) as the manufacturer instructed. The concentration of total RNAs solution was adjusted to 2mg/ml by RNase-free water. Then a small RNA (<200nt) separation buffer (2ml 50%(w/v) PEG 8000, 1ml 5M NaCl solution, 3ml RNase-free water) was prepared in a 15ml tube, 10mg (5ml) of mouse liver total RNAs were added to the tube and mixed with separation buffer by inverting the tube serval times. The tube was put on ice for 30mins followed by centrifuging at 12000rpm and 4 °C for 20mins. 9ml supernatant was collected and put into a new 50ml tube, following by adding 0.9ml NaAc solution (Invitrogen^TM^ AM9740), 29ml Ethanol, and 20ul Linear Acrylamide (Invitrogen^TM^ AM9520) to the tube. The tube was put in −20°C overnight and then centrifuged at 12000rpm and 4 °C for 20mins to get small RNAs (<200nt). Small RNA (<200nt) solution was adjusted to a concentration of 1mg/ml, then 1ml small RNA solution was transferred into a RNase-free 2ml tube followed by adding 6ul 100uM biotinylated tRNA-Glu DNA probe, 26ul 20x SSC solution (Invitrogen^TM^ 15557044) and 15ul RNase inhibitor (NEB M0314L). The 2ml tube was put in 50 °C water bath overnight. Transfer 200ul Streptavidin Sepharose (Cytiva 17511301) to a 1.5ml Ultrafree-MC tube (Millipore UFC30GV0S), then centrifuge at 2500g and room temperature (RT) for 30 seconds. Add 400 μL 20mM Tris-HCl (pH7.5) to the MC tube to wash Streptavidin Sepharose followed by centrifuging at 2500g and RT for 30 seconds. Repeat the washing process by 4 times. Next, transfer the hybridization solution to the MC tube to mix with the Streptavidin Sepharose, then transfer the solution and Streptavidin Sepharose back to the 2ml tube and put the tube on the shaker for 30 mins at RT. Then transfer the hybridization solution and Streptavidin Sepharose back to the MC tube, centrifuge at 2500g, and RT for 1 minute. Discard the flows and add 500ul 0.5X SSC to wash the Streptavidin Sepharose followed by centrifuging at 2500g and RT for 1 minute. Repeat the washing process by 5 times. Then add 500ul RNase-free water to the MC tube and incubate at 70°C for 15mins followed by centrifuging at 2500g and RT for 1mins to elute the RNAs which are complementary to the probe. Repeat the elution process by 3 times and a total of 1.5ml elution flows can be collected into a new 15ml tube. Then add 150ul NaAc solution, 4.5ml Ethanol and 10ul linear acrylamide to the elution flow, put in a tube in −20°C overnight and then centrifuge at 12000rpm and 4°C for 20mins to get RNAs which are complementary to the tRNA-Glu probe. Load these RNAs into the 7M Urea-PAGE gel to run electrophoresis, cut the main tRNA bind and perform the gel recovery to get enriched tRNA-Glu for MLC Seq. The enriched tRNA-Glu was also validated by the Northern Blot. The gel recovery process was described in our previous paper^7^. The probe was synthesized by IDT and the sequence was listed below: tRNA-Glu pulldown probe: 5’-Biotin-CTAACCACTAGACCACCAGGGA.

### Northern Blot

Northern Blot was performed as previously described^8^. RNA was separated by 10% Urea-PAGE gel stained with SYBR Gold, and immediately imaged, then transferred to positively charged Nylon membranes (Roche, 11417240001), and UV cross-linked with an energy of 0.12 J. Membranes were pre-hybridized with DIG Easy Hyb solution (Roche, REF: 11603558001) for 1h at 42 °C. To detect miRNAs, tsRNAs and rsRNAs in the total RNA and 15-50nt small RNAs, membranes were incubated overnight (12–16h) at 42 °C with DIG-labelled oligonucleotides probes synthesized by Integrated DNA Technologies (IDT). Then the membranes were washed twice with low stringent buffer (2× SSC with 0.1% (wt/vol) SDS) at 42 °C for 15 min each, then rinsed twice with high stringent buffer (0.1× SSC with 0.1% (wt/vol) SDS) for 5 min each, and then finally rinsed in washing buffer (1× SSC) for 10 min. Following the washes, the membranes were transferred into 1× blocking buffer (Roche, REF:11096176001) and incubated at room temperature for 3 h, after which the Anti-Digoxigenin-AP Fab fragments (Roche, REF: 11093274910) was added into the blocking buffer at a ratio of 1:10,000 and incubate for an additional half-hour at room temperature. The membranes were then washed four times with DIG washing buffer (1× maleic acid buffer, 0.3% Tween-20) for 15 min each, then incubated in DIG detection buffer (0.1 M Tris-HCl, 0.1 M NaCl, pH 9.5) for 5 min, and then coated with CSPD ready-to-use reagent (Roach REF: 11755633001) The membranes were incubated in the dark with the CSPD reagent for 30 min at 37 °C before imaging with ChemiDoc™ MP Imaging System (BIO-RAD). tRNA-Glu northern blot was synthesized by IDT and the sequence was listed below: tRNA-Glu Northern blot probe: 5’-DIG-CTAACCACTAGACCACCA.

### Treatment of tRNA-Glu with AlkB enzyme

The tRNA-Glu was incubated in 50 μl reaction mixture that contains 50 mM HEPES, pH 8.0 (Alfa Aesar, J63578), 75 μM ferrous ammonium sulfate (pH 5.0), 1 mM α-ketoglutaric acid (Sigma, K1128-25G), 2 mM sodium ascorbate, 50 μg/ml BSA (Sigma, A7906-500G), 2.5 μl RNase inhibitor (NEB M0314L), 200ng AlkB enzyme and 200 ng tRNA-Glu at 37°C for 30 mins (Recommended ratio of AlkB enzyme to RNA at 1:1 or above). Then the mixture was added into 500 μl TRIzol^TM^ reagent to perform the RNA isolation procedure as the manufacturer instructed.

## Data and Code Availability

All MS sequence datasets, pseudo and source codes for algorithms, and *de novo* MS sequence reads will be available upon request. They will be published in a peer-reviewed publication.

## Acknowledgments

The authors acknowledge grant support from the USA National Institutes of Health R21 HG009576 to S.Z. and W.L., R56 HG011099 to S.Z., R01 HD092431 and R01 ES032024 to Q.C., and R01 AI116812 and R21 AI113771to X.B.

