## Supplementary material for "MLC Seq: *De novo* sequencing of full-length tRNA isoforms by mass ladder complementation": SI

### SUPPORTING INFORMATION

**Table of Contents**


| Table of Contents | Page |
|---|---|
| Figure S 1 | 2 |
| Figure S 2 | 3 |
| Figure S 3 | 4 |
| Table S 1 | 5 |
| Table S 2 | 6 |

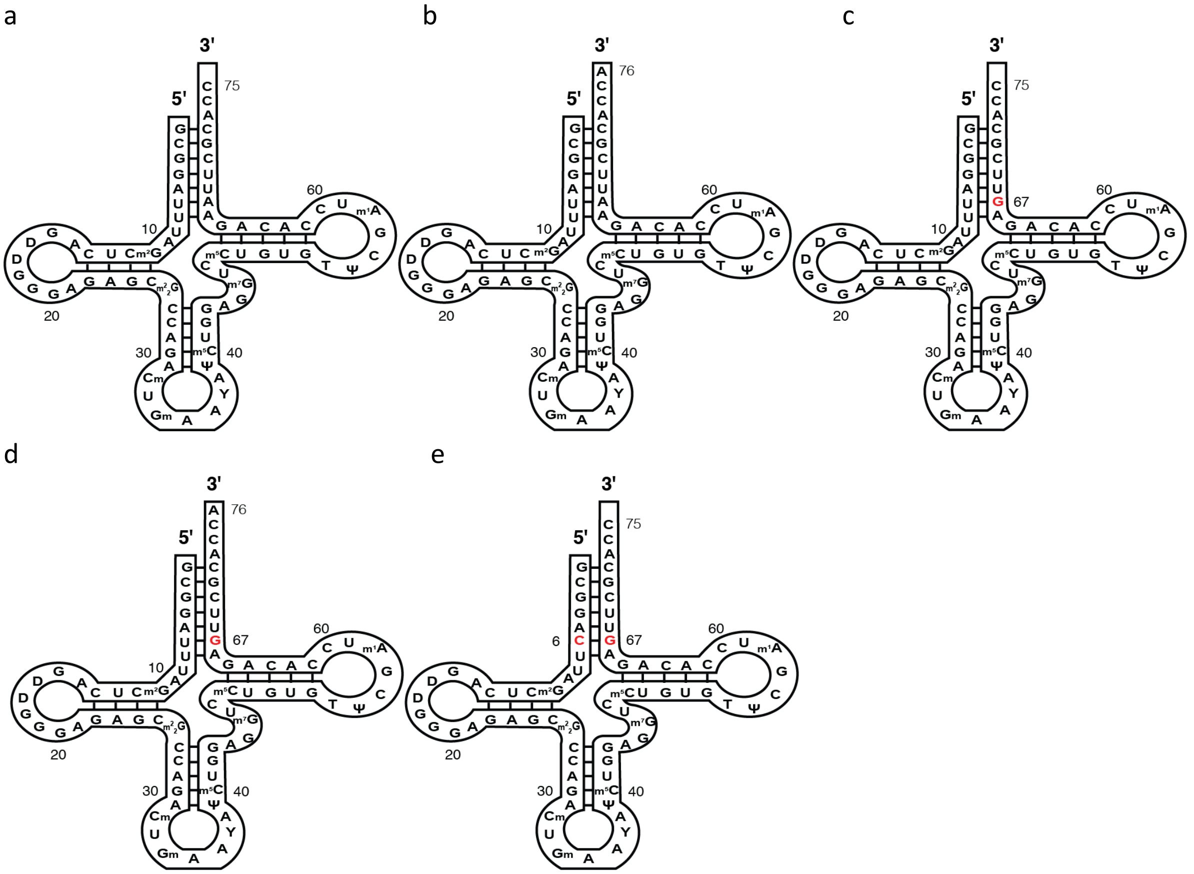


Figure S 1. MLC Seq results for 5 isoforms of tRNA-Phe, including (a) 75 nt tRNA-Phe with a CC-tail at the 3´-end (PheIF1). (b) 76 nt tRNA-Phe with a CCA tail at the 3´-end (PheIF2). (c) 75 nt tRNA-Phe +O (PheIF3). An A to G transition was observed at position 67. (d) 76 nt tRNA-Phe +O (PheIF4). An A to G transition was observed at position 67. (e) 75 nt tRNA-Phe plus 14.9373 Da (PheIF7). An A to G transition at position 67, and a U to C transition at position 6.

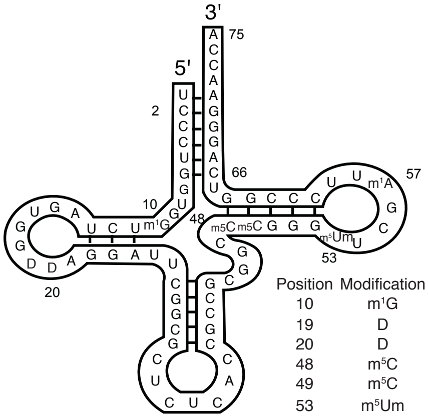


| Position | Modification | Ratio |
|---|---|---|
| 10 | $m^1G$ | 48% |
| 19 | D | 100% |
| 20 | D | 100% |
| 48 | $m^5C$ | 100% |
| 49 | $m^5C$ | 100% |
| 53 | $m^5Um$ | 100% |
| 57 | $m^1A$ | 68% |

Figure S 2. MLC Seq results of wild-type tRNA-Glu isoforms from mouse liver.

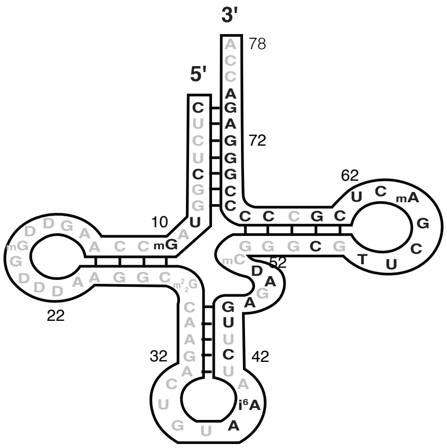


Figure S 3. MLC Seq of minor tRNA-Tyr (TyrIF1) in the yeast tRNA-Phe sample (Sigma).

Table S 1. tRNA isoforms from yeast tRNA-Phe and tRNA-Glu from mouse liver. Their names in this paper are listed here. (a) yeast tRNA-Phe. (b) tRNA-Glu extract from mouse liver. (c) tRNA-Glu extract from mouse liver treated with AlkB enzyme. (AD= Acid Degradation).

a

| Isoform Name | Mass (Before AD) | Mass (After AD) | Figure |
|---|---|---|---|
| **PheIF1** | 24610.4911 | 24252.3692 | Fig1, Fig3, Fig4 |
| **PheIF2** | 24939.5478 | 24581.3166 | Fig1, Fig4 |
| **PheIF3** | 24626.4639 | 24268.3038 | Fig2, Fig3, Fig4 |
| **PheIF4** | 24955.5224 | 24597.3533 | Fig2, Fig4 |
| **PheIF5** | | 24280.3055 | Fig4 |
| **PheIF6** | 24305.4031 | | Fig2 |
| **PheIF7** | | 24267.3065 | Fig S1 |
| **TyrIF1** | 25334.6284 | 25334.5688 | Fig2 |

b

| Isoform Name | Mass (Before AD) | Mass (After AD) | Figure |
|---|---|---|---|
| **GluIF1** | 24217.3144 | 24217.3031 | Fig5 |
| **GluIF2** | 24203.3106 | 24203.2936 | Fig5 |
| **GluIF3** | 24204.2858 | | Fig5 |
| **GluIF4** | 24189.3075 | 24189.2970 | Fig5 |
| **GluIF5** | 24218.3278 | 24218.3091 | Fig5 |
| **GluIF6** | | | |
| **GluIF7** | 24232.2884 | 24232.2913 | Fig5 |

c

| Isoform Name | Mass (Before AD) | Mass (After AD) | Figure |
|---|---|---|---|
| **GluIF1** | 24217.2662 | | Fig5 |
| **GluIF2** | 24203.3083 | 24203.2847 | Fig5 |
| **GluIF3** | 24204.2808 | 24204.2918 | Fig5 |
| **GluIF4** | 24189.2929 | 24189.2775 | Fig5 |
| **GluIF5** | 24218.2581 | 24218.2786 | Fig5 |
| **GluIF6** | 24219.2609 | 24219.2739 | Fig5 |
| **GluIF7** | | | |

Table S 2. Monoisotopic masses and intensities of ladder fragments from tRNA-Glu isoforms GluIF2 and GluIF4 before and after AlkB treatment.

| | Before AlkB | | | | After AlkB | | | |
|---|---|---|---|---|---|---|---|---|
| | LadderA(GluIF2) | | LadderB(GluIF4) | | LadderA(GluIF2) | | LadderB(GluIF4) | |
| **position** | **Mass** | **Intensity** | **Mass** | **Intensity** | **Mass** | **Intensity** | **Mass** | **Intensity** |
| **1** | 24203.2936 | 8.89E+05 | 24189.2970 | 1.59E+05 | 24203.2800 | 2.24E+06 | 24189.2775 | 3.64E+05 |
| **2** | | | | | 23817.3072 | 1.20E+04 | | |
| **3** | | | | | 23512.2674 | 2.18E+04 | | |
| **4** | | | | | 23207.2289 | 2.01E+04 | | |
| **5** | | | | | 22902.1913 | 7.77E+03 | 22888.1771 | 6.92E+02 |
| **6** | | | | | 22596.1385 | 1.62E+04 | | |
| **7** | 22251.1192 | 9.12E+02 | | | 22251.1090 | 2.14E+03 | 22237.0997 | 2.14E+04 |
| **8** | | | | | 21906.0558 | 2.79E+03 | 21892.0397 | 1.87E+04 |
| **9** | | | | | 21600.0380 | 6.68E+03 | 21586.0332 | 2.30E+04 |
| **10** | 21254.9766 | 7.91E+02 | 21240.9579 | 8.60E+02 | 21254.9684 | 2.54E+03 | 21240.9642 | 3.98E+04 |
| **11** | | | | | 20895.9066 | 2.26E+04 | | |
| **12** | | | | | 20589.8996 | 4.72E+04 | 20575.8908 | 6.15E+02 |
| **13** | | | | | 20284.8135 | 7.12E+04 | 20270.8364 | 1.03E+03 |
| **14** | | | | | 19978.8166 | 1.36E+05 | 19964.8073 | 2.22E+03 |
| **15** | | | | | 19649.7750 | 3.97E+04 | | |
| **16** | 19304.7221 | 8.51E+02 | | | 19304.7469 | 4.83E+04 | | |
| **17** | | | | | 18998.6964 | 6.52E+04 | | |
| **18** | 18653.6365 | 1.79E+03 | | | 18653.6479 | 8.46E+04 | | |
| **19** | | | | | 18308.6069 | 8.04E+04 | 18294.5805 | 8.04E+02 |
| **20** | 18000.5384 | 3.04E+03 | | | 18000.5515 | 9.94E+04 | | |
| **21** | 17692.4951 | 3.37E+03 | | | 17692.5142 | 1.17E+05 | 17678.5020 | 1.48E+03 |
| **22** | 17363.4611 | 8.03E+02 | | | 17363.4669 | 7.22E+04 | | |
| **23** | | | | | 17018.4063 | 8.04E+04 | 17004.3860 | 1.19E+03 |
| **24** | 16673.3450 | 3.74E+03 | | | 16673.3566 | 1.27E+05 | 16659.3435 | 1.66E+03 |
| **25** | 16344.3003 | 1.55E+03 | | | 16344.2982 | 8.11E+04 | | |
| **26** | 16038.2746 | 5.66E+03 | | | 16038.2691 | 1.25E+05 | 16024.2529 | 4.76E+03 |
| **27** | 15732.2568 | 1.31E+03 | | | 15732.2427 | 9.79E+04 | | |
| **28** | 15427.2039 | 2.23E+03 | | | 15427.2036 | 9.38E+04 | 15413.1733 | 1.80E+03 |
| **29** | 15082.1376 | 1.31E+03 | | | 15082.1563 | 9.72E+04 | 15068.1361 | 1.67E+03 |
| **30** | 14737.1017 | 9.71E+02 | | | 14737.1094 | 6.25E+04 | | |
| **31** | 14432.0729 | 1.40E+03 | | | 14432.0742 | 9.39E+04 | 14418.0397 | 1.89E+03 |
| **32** | 14087.0200 | 2.45E+03 | | | 14087.0251 | 8.13E+04 | | |
| **33** | 13781.9715 | 2.96E+03 | | | 13781.9750 | 1.08E+05 | 13767.9573 | 1.85E+03 |
| **34** | 13475.9240 | 4.11E+03 | 13461.9259 | 2.63E+02 | 13475.9362 | 1.47E+05 | 13461.9225 | 6.13E+03 |
| **35** | 13170.9032 | 2.93E+03 | | | 13170.8950 | 1.13E+05 | 13156.8537 | 6.47E+03 |
| **36** | 12864.8734 | 3.39E+03 | | | 12864.8804 | 1.32E+05 | 12850.8011 | 7.28E+03 |
| **37** | 12559.8332 | 5.85E+03 | | | 12559.8322 | 1.70E+05 | 12545.8006 | 7.04E+03 |
| **38** | 12230.7758 | 3.61E+03 | | | 12230.7639 | 8.55E+04 | 12216.7525 | 2.23E+03 |
| **39** | 11925.7393 | 3.09E+03 | | | 11925.7069 | 9.92E+04 | 11911.6726 | 1.97E+03 |
| **40** | 11620.6897 | 3.98E+03 | | | 11620.6619 | 1.06E+05 | | |
| **41** | 11275.6166 | 9.71E+02 | | | 11275.6314 | 9.30E+04 | 11261.6174 | 1.50E+03 |
| **42** | 10970.6036 | 2.77E+03 | | | 10970.5926 | 8.89E+04 | | |
| **43** | 10665.5554 | 3.33E+03 | | | 10665.5516 | 1.28E+05 | 10651.5421 | 1.10E+03 |
| **44** | 10320.4971 | 9.34E+02 | | | 10320.4975 | 8.01E+04 | | |
| **45** | 10015.4680 | 3.87E+03 | | | 10015.4713 | 1.04E+05 | 10001.4557 | 1.26E+03 |
| **46** | 9670.4181 | 4.56E+03 | | | 9670.4143 | 1.14E+05 | 9656.3829 | 2.28E+03 |
| **47** | 9325.3617 | 3.36E+03 | | | 9325.3648 | 1.01E+05 | 9311.3207 | 1.73E+03 |
| **48** | 9020.3210 | 5.42E+03 | | | 9020.3260 | 1.93E+05 | 9006.2904 | 4.05E+03 |
| **49** | 8701.2693 | 9.50E+03 | | | 8701.2692 | 2.04E+05 | 8687.2462 | 4.60E+03 |
| **50** | 8382.2106 | 5.77E+03 | | | 8382.2071 | 1.87E+05 | 8368.1957 | 2.36E+03 |
| **51** | 8037.1626 | 5.92E+03 | | | 8037.1638 | 1.45E+05 | 8023.1421 | 1.51E+03 |
| **52** | 7692.1167 | 7.63E+03 | | | 7692.1117 | 1.06E+05 | | |
| **53** | 7347.0403 | 3.38E+03 | | | 7347.0568 | 8.23E+04 | | |
| **54** | | | | | | | | |
| **55** | 6706.9878 | 1.18E+04 | | | 6706.984006 | 243700.88, 182342.89 | | |
| **56** | | | | | | | | |
| **57** | 6056.8913 | 8.93E+03 | | | 6056.8916 | 9.00E+04 | | |
| **58** | 5727.8385 | 3.31E+04 | | | 5727.8383 | 1.66E+04 | | |
| **59** | 5421.8146 | 4.15E+04 | | | 5421.8173 | 1.30E+04 | | |
| **60** | 4810.746709 | 37734.18, 37402.21 | | | 5115.788530 | 25765.42, 19640.6 | | |
| **61** | 4505.7039 | 1.60E+04 | | | | | | |
| **62** | 4200.6641 | 1.79E+04 | | | | | | |
| **63** | 3855.6107 | 5.51E+03 | | | | | | |
| **64** | | | | | | | | |
| **65** | | | | | | | | |
| **66** | | | | | | | | |
| **67** | 2570.3924 | 1.58E+04 | | | | | | |
| **68** | | | | | | | | |
| **69** | | | | | | | | |
| **70** | | | | | | | | |
| **71** | | | | | | | | |
| **72** | | | | | | | | |
| **73** | | | | | | | | |
| **74** | | | | | | | | |
| **75** | | | | | | | | |
